# Exploring Deep Physiological Models for Nociceptive Pain Recognition

**DOI:** 10.1101/622431

**Authors:** Patrick Thiam, Peter Bellmann, Hans A. Kestler, Friedhelm Schwenker

## Abstract

Standard feature engineering involves manually designing and assessing measurable descriptors based on some expert knowledge in the domain of application, followed by the selection of the best performing set of designed features in order to optimize an inference model. Several studies have shown that this whole manual process can be efficiently replaced by deep learning approaches which are characterized by the integration of feature engineering, feature selection and inference model optimization into a single learning process. Such techniques have proven to be very successful in the domain of image processing and have been able to attain state-of-the-art performances while significantly outperforming traditional approaches based on hand-crafted features. In the following work, we explore deep learning approaches for the analysis of physiological signals. More precisely, deep learning architectures are designed for the assessment of measurable physiological channels in order to perform an accurate classification of different levels of artificially induced nociceptive pain. Most of the previous works related to pain intensity classification based on physiological signals rely on a carefully designed set of hand-crafted features in order to achieve a relatively good classification performance. Therefore, the current work aims at building competitive pain intensity classification models without the need of domain specific expert knowledge for the generation of relevant features. The assessment of the designed deep learning architectures is based on the *BioVid Heat Pain Database (Part A)* and experimental validation demonstrates that the proposed uni-modal architecture for the electrodermal activity (EDA) and the deep fusion approaches significantly outperform previous classification methods reported in the literature, with respective average performances of 85.03% and 83.76% for the binary classification experiment consisting of the discrimination between the baseline level and the pain tolerance level (*T*_0_*vs*.*T*_4_) in a *Leave*-*One*-*Subject*-*Out* (LOSO) cross-validation evaluation setting.

## Introduction

Conventional machine learning approaches are built upon a set of carefully engineered representations, which consist of measurable parameters extracted from raw data. Based on some expert knowledge in the domain of application, a feature extractor is designed and used to extract relevant information in the form of a feature vector from the preprocessed raw data. This high level representation of the input data is subsequently used to optimise an inference model. Although such approaches have proven to be very effective and can potentially lead to state-of-the-art results (given that the set of extracted descriptors is suitable for the task at hand), the corresponding performance and generalisation capability is limited by the reliance on expert knowledge as well as the inability of the designed model to process raw data directly and to dynamically adapt to related new tasks.

Meanwhile, deep learning approaches [1] automatically generate suitable representations by applying a succession of simple and non-linear transformations on the raw data. A deep learning architecture consists of an hierarchical construct of several processing layers. Each processing layer is characterised by a set of parameters that are used to transform its input (which is the representation generated by the previous layer) into a new and more abstract representation. This specific hierarchical combination of several non-linear transformations enables deep learning architectures to learn very complex functions as well as abstract descriptive (or discriminative) representations directly from raw data [2]. Moreover, the hierarchical construct characterising deep learning architectures offers more flexibility when it comes to adapt such approaches to new and related tasks. Hence, deep learning approaches have been outperforming previous state-of-the-art machine learning approaches, especially in the field of image processing [3–7]. Similar performances have been achieved in the field of speech recognition [8, 9] and natural language processing [10, 11].

A steadily growing amount of work has been exploring the application of deep learning approaches on physiological signals. Martinéz et al. [12] were able to significantly outperform standard approaches built upon hand-crafted features by using a deep learning algorithm for affect modelling based on physiological signals (two physiological signals consisting of Skin Conductance (SC) and Blood Volume Pulse (BVP) were used in this specific work). The designed approach consisted of a multi-layer Convolutional Neural Network (CNN) [13] combined with a single-layer perceptron (SLP). The parameters of the CNN were trained in an unsupervised manner using denoising auto-encoders [14]. The SLP was subsequently trained in a supervised manner using backpropagation [15] to map the outputs of the CNN to the target affective states. In [16], the authors proposed a multiple-fusion-layer based ensemble classifier of stacked auto-encoder (MESAE) for emotion recognition based on physiological data. A physiological-data-driven approach was proposed in order to identify the structure of the ensemble. The architecture was able to significantly outperform the existing state-of-the-art performance. A deep CNN was also successfully applied in [17] for arousal and valence classification based on both electrocardiogram (ECG) and Galvanic Skin Response (GSR) signals. In [18], a hybrid approach using CNN and Long Short-Term Memory (LSTM) [19] Recurrent Neural Network (RNN) was designed in order to automatically extract and merge relevant information from several data streams stemming from different modalities (physiological signals, environmental and location data) for emotion classification. Moreover, deep learning approaches have been applied on electromyogram (EMG) signals for gesture recognition [20, 21] or hand movement classification [22, 23]. Most of the reported approaches consist of first transforming the processed EMG signal into a two dimensional (time-frequency) visual representation (such as a spectrogram or a scalogram) and subsequently using a deep CNN architecture to proceed with the classification. A similar procedure has been used in [24] for the analysis of electroencephalogram (EEG) signals. These are just some examples of an increasingly growing field of experimentation for deep learning architectures. A better overview of deep learning approaches applied to physiological signals can be found in [25] and [26].

The current work focuses on the application of deep learning approaches for nociceptive pain recognition based on physiological signals (EMG, ECG and electrodermal activity (EDA)). Several deep learning architectures are proposed for the assessment of measurable physiological parameters in order to perform an end-to-end classification of different levels of artificially induced nociceptive pain. The current work aims at achieving state-of-the-art classification performances based on feature learning, therefore removing the reliance on expert knowledge for the optimization of reliable pain intensity classification models, since most of the previous works on pain intensity classification involving autonomic parameters rely on a carefully designed set of hand-crafted features. Most recently, Thiam et al. [27] provided the results for a row of pain intensity classification experiments based on the *SenseEmotion Database* (SEDB) [28], by using several fusion architectures to merge hand-crafted features extracted from different input modalities, including physiological, audio and video channels. Thereby, the combination of the features extracted from the recorded signals was compared for different fusion approaches, i.e. the fusion at feature level, the fusion at the classifiers’ output level and the fusion at an intermediate level. Random Forests [29] were used as the base classifiers. In [30], Kessler et al. combined camera photoplethysmography input signals with ECG and EMG signals in order to proceed with a user-independent pain intensity classification using the same dataset. The authors used a fusion architecture at feature level with Random Forests and Support Vector Machines (SVM) [31] as base classifiers.

In [32–34], the authors perform different pain intensity classification experiments based on the *BioVid Heat Pain Database* [35] (BVDB). All the conducted experiments are based upon a carefully selected set of features extracted from both physiological and video channels. The classification is also performed using either Random Forests or SVMs. In [36], Kächele et al. perform a user-independent pain intensity classification evaluation based on physiological input signals, using the same dataset. The authors used the whole data from all recorded pain levels in a classification, as well as a regression setting with Random Forests as the base classification and regression models, respectively. Several personalisation techniques were designed and validated, based on meta information from the test subjects, distance measures and machine learning techniques. The same authors proposed an adaptive confidence learning approach for personalized pain estimation in [37] based on both physiological and video modalities. Thereby, the authors applied the fusion at feature level. The whole pain intensity estimation task was analysed as a regression problem. Random Forests were used as the base regression models. Moreover, a multi-layer perceptron (MLP) was applied to compute the confidence for an additional personalisation step. One of the recent works, including the physiological signals of both data sets (SEDB and BVDB) is [38]. The authors analysed different fusion approaches with fixed aggregating rules based on their merging level for the person-independent multi-class scenario using all available pain levels. Thereby, three of the most popular decision tree based classifier systems, i.e. Bagging [39], Boosting [40] and Random Forests were compared.

The remainder of the work is organised as follows. The proposed deep learning approaches as well as the dataset used for the validation of the approaches are described in the section Materials and Methods. Subsequently, a description of the results corresponding to the conducted assessments specific to each presented approach is provided in the section Results. Finally, the findings of the conducted experiments are discussed in the section Discussion and Conclusion, followed by the description of potential future works and a conclusion.

## Materials and Methods

### BioVid Heat Pain Database (BVDB)

The *BioVid Heat Pain Database* [35] (BVDB) was collected at Ulm University. It includes multi modal data recordings from healthy subjects exposed to different artificially induced pain stimuli under strictly controlled conditions. The pain elicitation in the form of heat was conducted through the professionally designed PATHWAY (http://www.medoc-web/products/pathway) thermode attached to the participants’ right forearm. Before the data was recorded, a personalised calibration step was undertaken for each participant to determine individual levels for the pain threshold, as well as the tolerance threshold. Therefore, starting at a temperature of 32°*C* (global pain free level *T*_0_ for all participants), the temperature was slowly increased until first, the participant felt a change from heat to pain (pain threshold *T*_1_), and second, the pain became hardly bearable (tolerance threshold *T*_4_). In addition, two in-between pain elicitation levels *T*_2_ and *T*_3_ were calculated, making the four individual pain levels *T*_1_, *T*_2_, *T*_3_, *T*_4_ equidistant. After the initial calibration steps, starting at the baseline temperature *T*_0_, each of the four individual pain levels was applied randomly 20 times for stimulating the participant. Each of the pain levels was held for a total of 4 seconds (sec). Each pain stimulation was followed by a rest period during which the baseline temperature was held for a random duration of 8 to 12 sec. Ninety subjects were recruited for the experiments. The participants covered three age groups, i.e. 18-35 years, 36-50 years and 51-65 years. Each group was equally distributed, including 15 male and 15 female subjects. In the current work, the designed approaches are assessed on the *BioVid Heat Pain Database (Part A)* since most of the related works were conducted based on this specific database. The database is publicly available and consists of a total of 87 participants. The second part of the BVDB dataset (Part B) consists of a total of 86 participants. The difference between the two parts apart from the number of participants is that in part B, additional physiological data were recorded (zygomaticus and corrugator EMG data).

During the experiments, three different physiological signals were recorded, namely electrodermal activity (EDA), eletrocardiogram (ECG) and electromyogram (EMG) (a sample of the recorded physiological signals is depicted in Fig 1). The EDA is an indicator for the skin conductance level and was measured at both, the participants’ index and ring fingers. The ECG signals measure the participants’ heart activity, like the heart rate, the interbeat interval and the heart rate variability. The EMG signal is an indicator for the muscle activity. The EMG signal of the current dataset consists of the muscle activities of the trapezius muscles, which are located at the back, in the shoulder area. In addition to the biopotentials, different video signals were recorded. Since we are only considering the physiological signals here, interested readers are referred to [35] to get further details on the whole data set. Having 20 elicitations for each level of pain elicitation every subject is represented by 100 sequences of numerical data points (time series).

**Fig 1.**
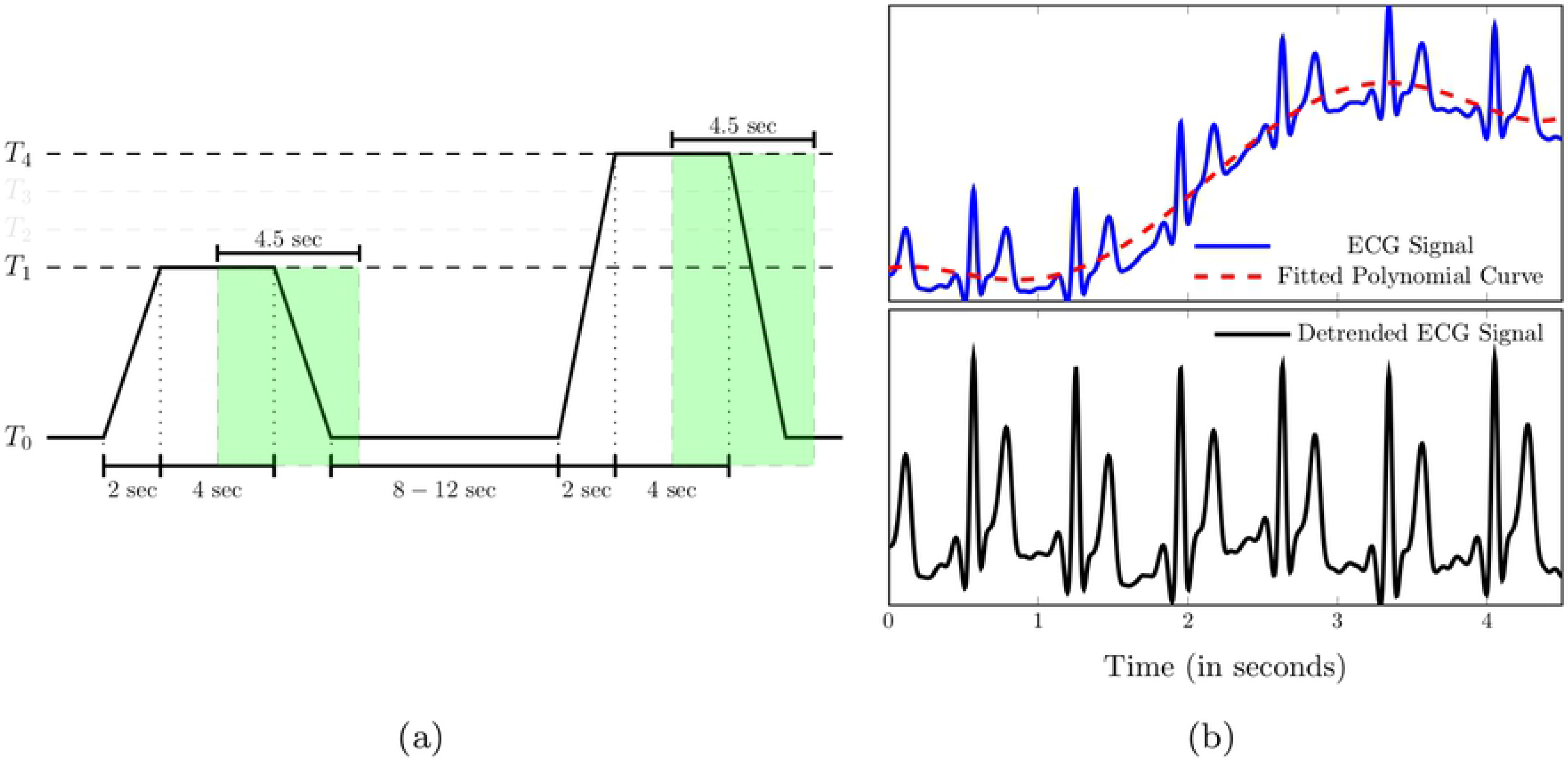
Recorded physiological data. From top to bottom: series of artificially induced pain elicitation (*T*_1_: pain threshold temperature, *T*_2_: first intermediate elicitation temperature, *T*_3_: second intermediate elicitation temperature, *T*_4_: pain tolerance temperature); EDA (*μ*S); EMG (*μ*V); ECG (*μ*V).

### Data Preprocessing

Prior to the classification experiments, the sampling rate of the recorded physiological modalities was reduced to 256 Hz, in order to reduce the computational requirements. Subsequently, the amount of noise and artefacts within the recorded data was significantly reduced by applying different signal preprocessing techniques on each specific modality. A third order low-pass Butterworth filter with a cut-off frequency of 0.2 Hz was applied on the EDA signal. The EMG signal was filtered by applying a fourth order bandpass Butterworth filter with a frequency range of [20, 250] Hz. Finally, a third order bandpass Butterworth filter with a frequency range of [0.1, 250] Hz was applied on the ECG signal. Furthermore, the data is segmented as proposed in [33], but rather than using 5.5 sec windows with a shift of 3 sec from the elicitations’ onset, the preprocessed signals were segmented into windows of length 4.5 sec, with a shift from the elicitations’ onset of 4 sec (see Fig 2(a)) based on the data driven signal segmentation approach that was recently proposed in [27]. Each signal extracted within this window constitutes a 1-D array of size 4.5 × 256 = 1152 and is later on used in combination with the corresponding level of pain elicitation to optimize and assess the designed deep classification architectures. Thus, each physiological modality specific to each single participant is represented by a tensor with the dimensionality 100 × 1152 × 1. Additionally, the segmented ECG signals were detrended by subtracting a 5^*th*^ degree polynomial least-squares fit from the filtered signal. This step was carried out to remove the artefacts stemming from the recorded EDA signal which could potentially bias the classification performance of the corresponding deep classification model (see Fig 2(b)). Finally, data augmentation was performed by shifting the 4.5 sec window of segmentation backward and forward in time with small shifts of length 250 milliseconds (ms) and a maximal total window shift of 1 sec in each direction, starting from the initial position of the window depicted in Fig 2(a). The signals extracted within these windows were subsequently used as training material for the optimization of the classification architectures.

**Fig 2.**
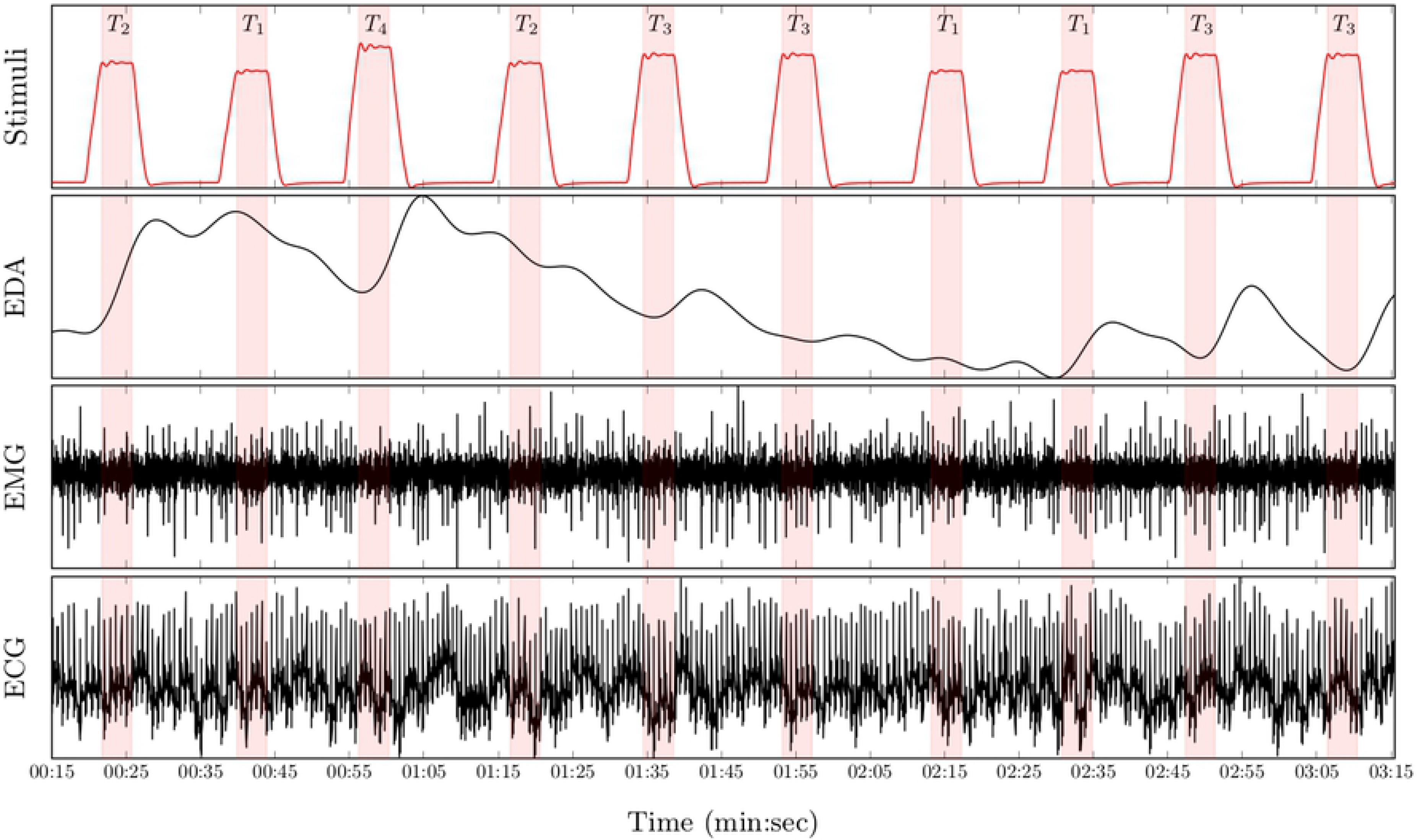
Data preprocessing. (a) Signal Segmentation. The classification experiments are performed on windows of length 4.5 sec with a temporal shift of 4 sec from the elicitations’ onset. (b) The ECG signal is further detrendet by subtracting a least-squares polynomial fit from the preprocessed signal.

### Uni-modal Deep Model Description

As mentioned earlier, the goal of the current work is to apply feature learning in order to alleviate the reliance on domain specific expert knowledge that occurs when relevant and adequate features are to be manually designed (hand-crafted features) in order to achieve state-of-the-art classification performances. Therefore, multi-layer CNNs are designed and fed with the preprocessed physiological signals in order to automatically compute relevant signal representations and at the same time optimize the classification architectures. In the following sections, *c* depicts the number of classes of the classification task.

CNNs [41, 42] constitute a distinct category of biologically inspired neural networks, which are characterised by a hierarchical structure of several processing layers. The input to a CNN is sequentially and progressively transformed by each specific layer and the back-propagated information stemming from the error computed between the network’s output and the expected output (ground-truth) is used to optimize the whole structure of the architecture in order to efficiently and effectively solve a classification or regression task. The basic processing layers of CNNs are *convolutional layers*, *pooling layers* and *fully connected layers*. *Convolutional layers* are characterised by a set of neurons (or kernels), whereby each specific neuron extracts a specific pattern of information from a patch of the layer’s input. Each neuron consists of a set of trainable weights, which size is determined by the patch’s size (or kernel size). The output of each neuron is calculated by applying a non-linear activation function (e.g. sigmoid function) on the weighted sum of the neuron’s input. Each neuron scans the layer’s input sequentially and the aggregation of the resulting local information extracted at each specific patch constitutes a feature map. Thus the output of a convolutional layer is a set of feature maps generated by the convolution of each neuron across the layer’s input. *Pooling layers* reduce the spatial resolution of the generated feature maps by merging semantically similar features. *Max Pooling* is a commonly used pooling approach and consists of computing the maximum value of a defined local patch (the size of the patch related to a specific pooling layer is referred in the current work as “pool size”) of each feature map. *Fully connected layers* are basically single-layer feed-forward networks that perform the classification or regression task based on the learned deep representations.

Several challenges emerge when it comes to optimize such architectures. One of those challenges is the so called *vanishing* or *exploding gradients* problem which is caused by the *internal covariate shift* (constant fluctuations in layers’ input distributions) occurring in deep architectures during the training process. In [43], the authors proposed a technique called *Batch Normalisation* to address this specific issue. Batch Normalisation consists of automatically learning the optimal scaling and shifting parameters of each layer’s input, so that each layer’s input is dynamically normalized, thus significantly reducing the effects of the internal covariate shift and stabilizing the training process. Another common challenge occurring when training CNNs is the *overfitting* problem caused by the large amount of parameters that have to be consistently and effectively optimized. Applying regularisation techniques can help to significantly reduce this issue. The authors in [44] introduced the *dropout* approach, which is one of the most commonly used regularisation technique for deep learning architectures. The *dropout* approach consists of randomly and temporarily removing a set of neurons (or units) from the neural network during each training step, each neuron having a fixed probability *p* ∈ [0, 1] to be retained. The resulting model is therefore more robust against overfitting and generalises better.

In the current work, the designed architectures are regularised using both techniques and the dropout rate is fixed to 25%. Moreover, the *Exponential Linear Unit* (ELU) function [45] defined in Eq. (1)

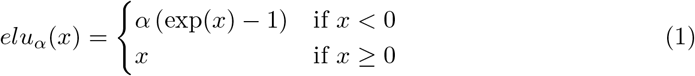

is used as activation function for both convolutional layers and fully connected layers (with *α* = 1), except for the last fully connected layer of each architecture where a *softmax* function defined in Eq. (2)

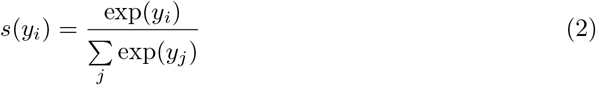

is used as activation function, with 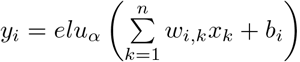 (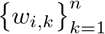 represents the set of weights of the *i*^*th*^ neuron, *b*_*i*_ represents the bias term of the *i*^*th*^ neuron and *x* = (*x*_1_,…, *x*_*i*_,…, *x*_*n*_) represents the output of the precedent fully connected layer). The designed architectures for each physiological signal are based on 1-D convolutional layers and are described in Table 1.

**Table 1.**
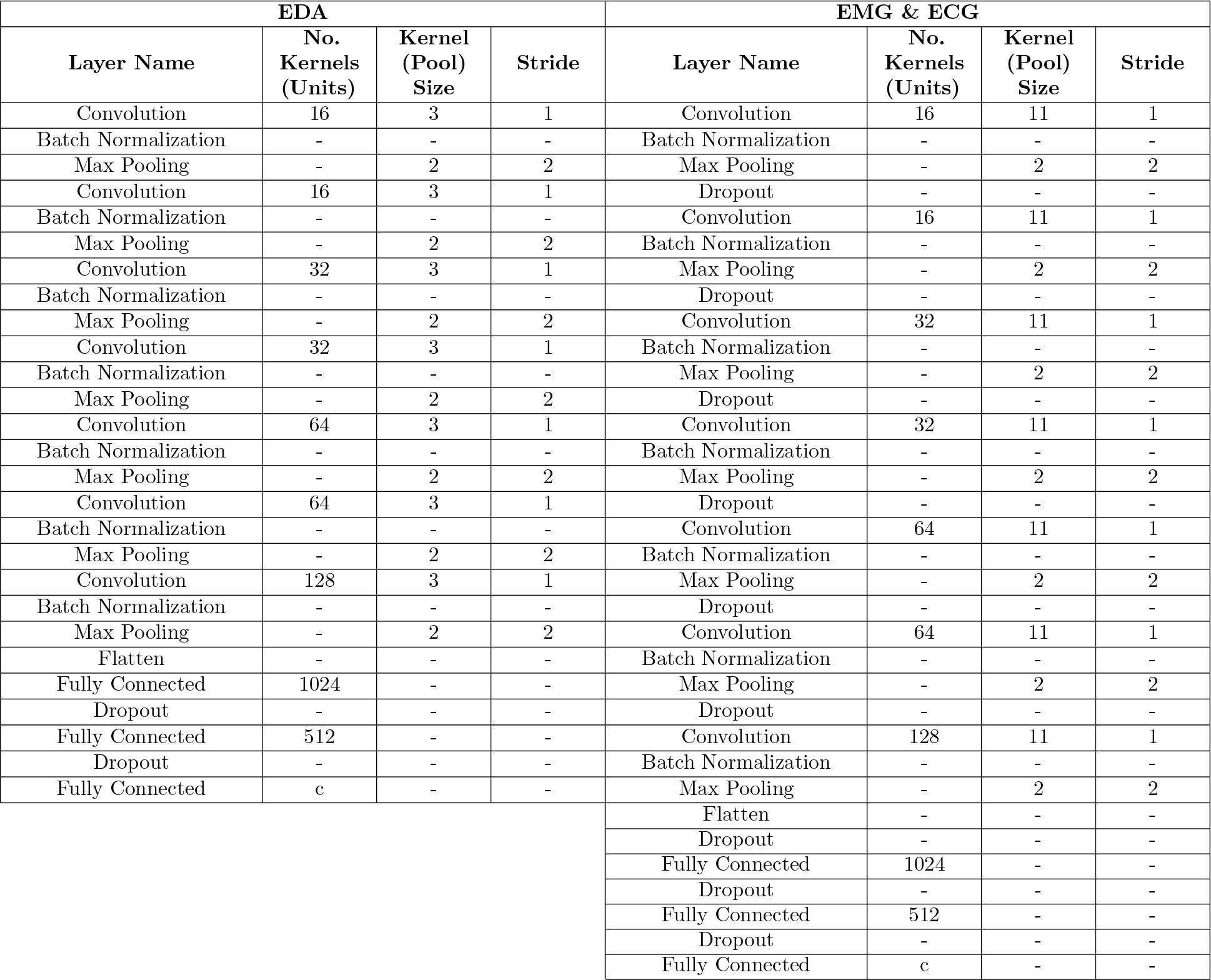
Deep classification architectures for each of the recorded physiological modality. ELU is used as activation function in both convolutional and fully connected layers, except for the last fully connected layer where a *softmax* activation function is used. The networks are further regularized by using *dropout* layers with a fixed dropout rate of 25%.

| EDA |  |  |  | EMG & ECG |  |  |  |
| --- | --- | --- | --- | --- | --- | --- | --- |
| Layer Name | No. Kernels (Units) | Kernel (Pool) Size | Stride | Layer Name | No. Kernels (Units) | Kernel (Pool) Size | Stride |
| Convolution | 16 | 3 | 1 | Convolution | 16 | 11 | 1 |
| Batch Normalization | - | - | - | Batch Normalization | - | - | - |
| Max Pooling | - | 2 | 2 | Max Pooling | - | 2 | 2 |
| Convolution | 16 | 3 | 1 | Dropout | - | - | - |
| Batch Normalization | - | - | - | Convolution | 16 | 11 | 1 |
| Max Pooling | - | 2 | 2 | Batch Normalization | - | - | - |
| Convolution | 32 | 3 | 1 | Max Pooling | - | 2 | 2 |
| Batch Normalization | - | - | - | Dropout | - | - | - |
| Max Pooling | - | 2 | 2 | Convolution | 32 | 11 | 1 |
| Convolution | 32 | 3 | 1 | Batch Normalization | - | - | - |
| Batch Normalization | - | - | - | Max Pooling | - | 2 | 2 |
| Max Pooling | - | 2 | 2 | Dropout | - | - | - |
| Convolution | 64 | 3 | 1 | Convolution | 32 | 11 | 1 |
| Batch Normalization | - | - | - | Batch Normalization | - | - | - |
| Max Pooling | - | 2 | 2 | Max Pooling | - | 2 | 2 |
| Convolution | 64 | 3 | 1 | Dropout | - | - | - |
| Batch Normalization | - | - | - | Convolution | 64 | 11 | 1 |
| Max Pooling | - | 2 | 2 | Batch Normalization | - | - | - |
| Convolution | 128 | 3 | 1 | Max Pooling | - | 2 | 2 |
| Batch Normalization | - | - | - | Dropout | - | - | - |
| Max Pooling | - | 2 | 2 | Convolution | 64 | 11 | 1 |
| Flatten | - | - | - | Batch Normalization | - | - | - |
| Fully Connected | 1024 | - | - | Max Pooling | - | 2 | 2 |
| Dropout | - | - | - | Dropout | - | - | - |
| Fully Connected | 512 | - | - | Convolution | 128 | 11 | 1 |
| Dropout | - | - | - | Batch Normalization | - | - | - |
| Fully Connected | c | - | - | Max Pooling | - | 2 | 2 |
|  |  |  |  | Flatten | - | - | - |
|  |  |  |  | Dropout | - | - | - |
|  |  |  |  | Fully Connected | 1024 | - | - |
|  |  |  |  | Dropout | - | - | - |
|  |  |  |  | Fully Connected | 512 | - | - |
|  |  |  |  | Dropout | - | - | - |
|  |  |  |  | Fully Connected | c | - | - |

### Multi-modal Deep Model Description

In order to further investigate the compatibility of the recorded physiological data, several fusion approaches based on CNNs are proposed. The information stemming from each modality is aggregated at different levels of abstraction.

The first approach depicted in Fig 3 consists of an early fusion method, where the aggregation is done at the lowest level of abstraction which consists of the preprocessed raw signals (input data). A 2-D representation of the input data is generated by concatenating the three physiological modalities along the temporal axis, resulting into a tensor with the dimensionality 3 × 1152 × 1. The resulting data is subsequently fed into a network consisting of 2-D convolutional layers. The motivation behind such an approach is to enable the architecture to dynamically learn an appropriate set of weights, which will generate feature maps consisting of relevant and compatible information extracted simultaneously from the recorded modalities, when applied to the 2-D data representation. The designed fusion architecture is described in Table 2.

Furthermore, two additional late fusion approaches are proposed (see Fig 4). Both approaches are based on the uni-modal CNN architectures described earlier (see section Uni-modal Deep Model Description). The first approach described in Fig 4(a) performs the aggregation of the information at the mid-level since it involves using intermediate representations of the input data. It consists of concatenating the outputs of the second fully connected layer of each modality specific architecture and feeding the resulting representation to an output layer with a *softmax* activation function.

**Fig 3.**
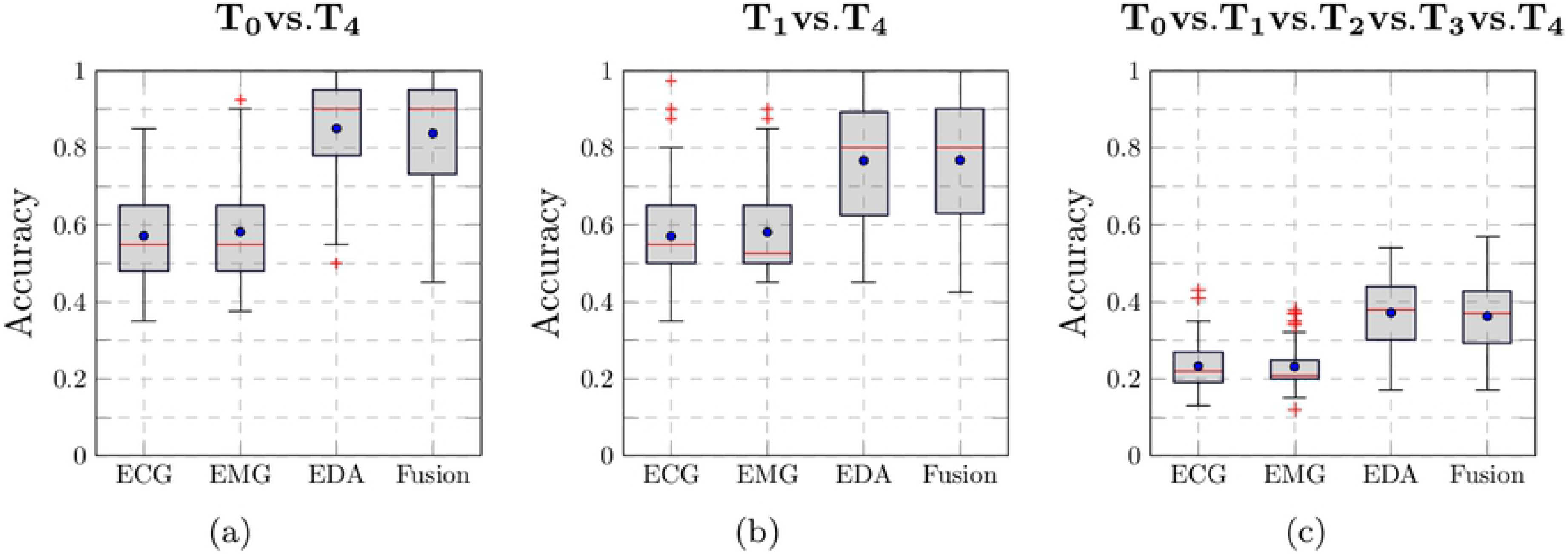
Early Fusion Architecture. A 2-D representation of the input data is generated by concatenating the three physiological modalities and is subsequently fed into the designed deep architecture.

**Table 2.** Early fusion deep CNN architecture. The architecture is based on 2-D convolutional layers. A 2-D representation of the input data is generated by concatenating the three physiological modalities resulting in a tensor with the dimensionality 3 × 1152 × 1. Similar to the previous architectures, ELU is used as activation function in both convolutional and fully connected layers, except for the last fully connected layer where a *softmax* activation function is used. The network is further regularized by using *dropout* layers with a fixed dropout rate of 25%.

| Layer Name | No. Kernels (Units) | Kernel (Pool) Size | Stride |
| --- | --- | --- | --- |
| Convolution | 16 | $2 \times 11$ | $1 \times 1$ |
| Convolution | 16 | $2 \times 11$ | $1 \times 1$ |
| Batch Normalization | - | - | - |
| Max Pooling | - | $1 \times 2$ | $1 \times 2$ |
| Dropout | - | - | - |
| Convolution | 32 | $1 \times 11$ | $1 \times 1$ |
| Batch Normalization | - | - | - |
| Max Pooling | - | $1 \times 2$ | $1 \times 2$ |
| Dropout | - | - | - |
| Convolution | 32 | $1 \times 11$ | $1 \times 1$ |
| Batch Normalization | - | - | - |
| Max Pooling | - | $1 \times 2$ | $1 \times 2$ |
| Dropout | - | - | - |
| Convolution | 64 | $1 \times 11$ | $1 \times 1$ |
| Batch Normalization | - | - | - |
| Max Pooling | - | $1 \times 2$ | $1 \times 2$ |
| Dropout | - | - | - |
| Convolution | 64 | $1 \times 11$ | $1 \times 1$ |
| Batch Normalization | - | - | - |
| Max Pooling | - | $1 \times 2$ | $1 \times 2$ |
| Flatten | - | - | - |
| Dropout | - | - | - |
| Fully Connected | 1024 | - | - |
| Dropout | - | - | - |
| Fully Connected | 512 | - | - |
| Dropout | - | - | - |
| Fully Connected | c | - | - |

**Fig 4.**
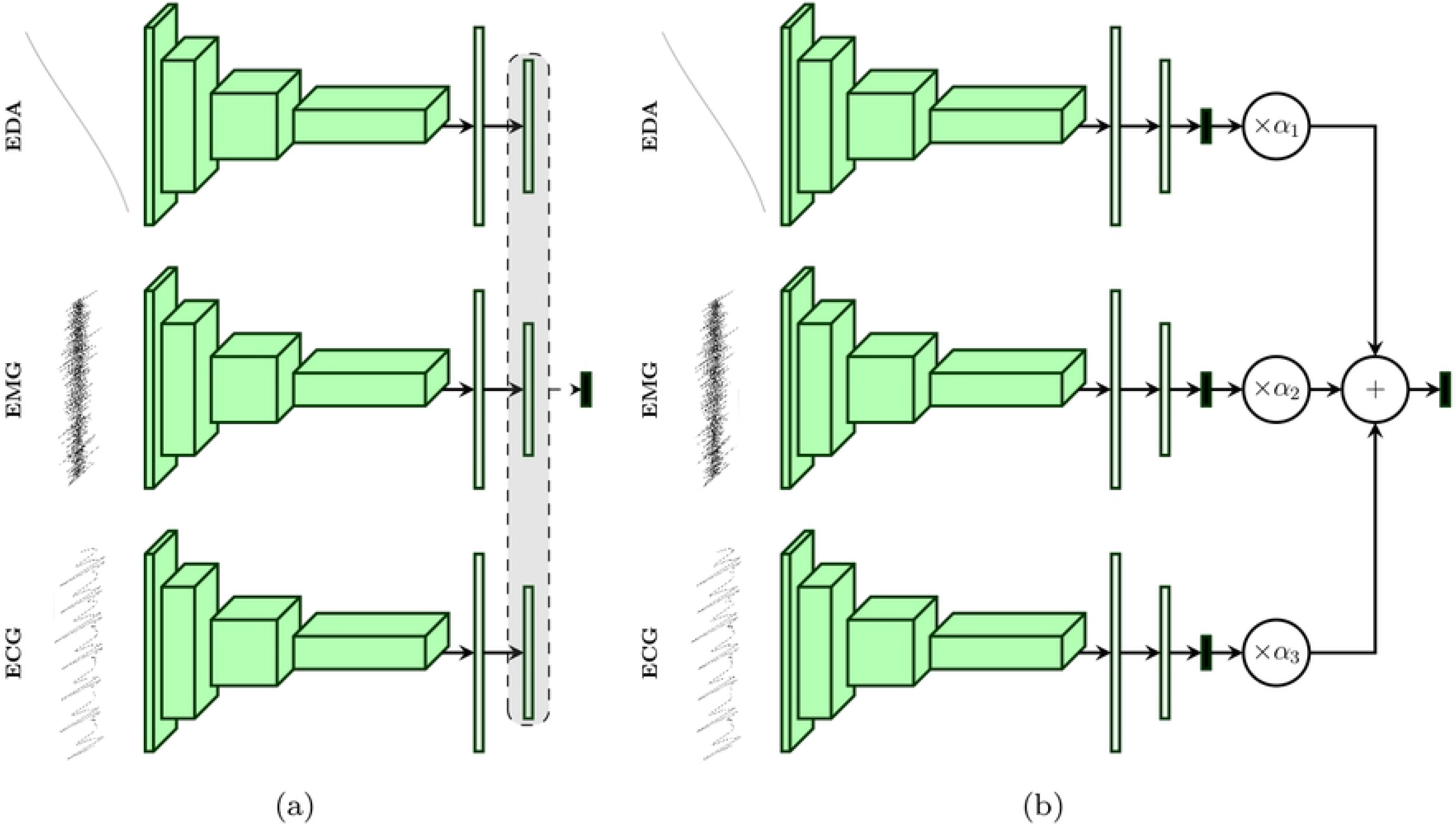
Late Fusion Architectures. (a) The features extracted by the second fully connected layer are concatenated and fed into the output layer. (b) A weighted average of the outputs of each uni-modal model is computed and fed into a softmax layer.

The second approach depicted in Fig 4(b) performs the aggregation at the highest level of abstraction, since it involves using the respective *softmax* layers’ outputs of each modality specific architecture. An additional layer consisting of a set of trainable positive parameters 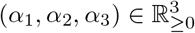 with a *softmax* activation function is directly connected to the outputs of each uni-modal architecture. For each modality specific architecture *i* ∈ {1, 2, 3} (since we are dealing with 3 physiological modalities), let {*θ*_*i,j*_ ∈ [0,1]: 1 ≤ *j* ≤ c} be the output values of the respective *softmax* layers. The output of the aggregation layer is computed by using the following formulas:

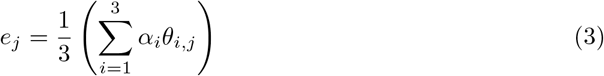

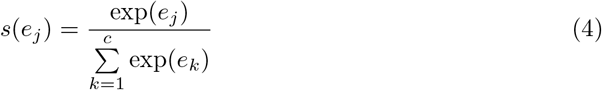

First a weighted average output of the class probabilities stemming from the uni-modal architectures is computed (see Eq. 3), and the corresponding class probabilities of the fusion architecture are subsequently deducted by applying a *softmax* function on the previously computed scores (see Eq. 4). Furthermore, the whole architecture is trained by using the loss function defined in Eq. 5.

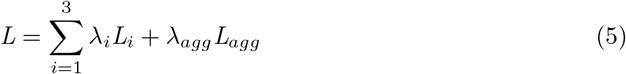

where *L*_1_, *L*_2_ and *L*_3_ are the loss functions of each modality specific architecture and *L*_*agg*_ is the loss function of the aggregation layer. The parameters *λ*_1_, *λ*_2_, *λ*_3_ and *λ*_*agg*_ are the corresponding weights for each of the loss functions. Once the architecture has been trained, unseen samples are classified based uniquely on the output of the aggregation layer. All described fusion approaches are subsequently trained in an *end*-*to*-*end* manner, which means that the fusion parameters are optimised at the same time as the parameters of each modality specific classification architecture. Furthermore, the parameters of each described architecture (uni-modal as well as multi-modal) are optimised using the cross entropy loss function defined in Eq. 6,

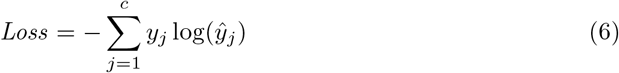

where *y*_*j*_ is the ground-truth value of the *j*^*th*^ class and 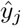 is the *j*^*th*^ output value of the *softmax* function. Concerning the second late fusion architecture, the cross entropy loss function is used for each uni-modal architecture as well as for the aggregation layer (*L*_1_ = *L*_2_ = *L*_3_ = *L*_*agg*_).

## Results

All previously described deep architectures are trained using the Adaptive Moment estimation (*Adam*) [46] optimisation algorithm with a fixed learning rate set empirically to 10^−5^. The training process consists of a total number of 100 epoches with the batch size set to 100. The weights of the loss function for the second late fusion architecture (see Fig 4(b)) are empirically set as follows: *λ*_1_ = *λ*_2_ = *λ*_3_ = 0.2, *λ*_*agg*_ = 0.4. The weight corresponding to the aggregation layer (*λ*_*agg*_) is set higher than the others in order to push the network to focus on the weighted combination of the single modality architectures’ outputs, and therefore to evaluate an optimal set of the weighting parameters {*α*_1_, *α*_2_, *α*_3_}. The implementation and evaluation of the described algorithms is done with the libraries Keras [47], Tensorflow [48] and Scikit-learn [49]. The evaluation of the architectures is performed in a *Leave*-*One*-*Subject*-*Out* (LOSO) cross-validation setting.

A performance evaluation of the designed architectures in a binary classification task consisting of the discrimination between the baseline temperature *T*_0_ and the pain tolerance temperature *T*_4_ is reported in Table 3. The achieved results based on CNNs are also compared to the state-of-the-art results reported in previous works. At a glance, the designed deep learning architectures outperform the state-of-the-art results in every setting, except for the ECG modality. Regarding the aggregation of all physiological modalities, the second late fusion architecture performs best and sets a new state-of-the-art fusion performance with an average accuracy of 83.76%, which even outperforms the best fusion results reported in [37], where the authors could achieve an average classification performance of 83.1% by using both physiological and video features. The deep architecture based on the EDA modality significantly outperforms all previously reported classification results with an average accuracy of 85.03%. Based on this finding, further binary classification experiments were conducted, based uniquely on the EDA modality and compared with similar results in previous works. The results depicted in Table 4 clearly show that the designed CNN architecture is able to consistently and significantly outperform previous approaches in all classification settings. EDA is also the best performing single modality, followed by EMG and ECG respectively, which is consistent with the results reported in previous works. Concerning the classification task *T*_0_*vs*.*T*_4_, even though the deep fusion architectures attain state-of-the-art fusion performances, they are unable to outperform the CNN architecture based uniquely on EDA. The aggregated information stemming from both modalities EMG and ECG seems to harm the optimisation process of the whole classification architecture in several cases (as can be seen in Fig 5(a)). It can also be seen in Fig 5(b), that the fusion architecture systematically assigned a higher weight value to EDA, while both ECG and EMG are assigned significantly lower weight values. EDA is assigned an average weight value of 0.29 with a standard deviation of 0.01 × 10^−2^. EMG is assigned an average weight value of 0.19 with a standard deviation of 1.67 × 10^−2^ and ECG is assigned an average weight value of 0.19 with a standard deviation of 1.26 × 10^−2^. Therefore, since the weighting parameters seem to improve the generalisation ability of the classification architecture, it is believed that an improved optimization of these specific parameters could further improve the performance of the designed system.

**Table 3.** Performance comparison to early work on the BVDB (Part A) for the classification task *T*_0_ vs.*T*_4_ in a LOSO cross-validation setting. The performance metric consists of the average accuracy (in %) over the LOSO cross-validation. The best performing approach for each modality and the aggregation of all modalities is depicted in bold.

| Method | ECG | EMG | EDA | Fusion |
| --- | --- | --- | --- | --- |
| Werner et al. [32] | **62.00** | 57.90 | 73.80 | 74.10 |
| Kächele et al. [36,37] | n.a. | n.a. | 81.10 | 82.73 |
| Our Approaches (CNNs) | 57.16 | **58.16** | **85.03** | Early Fusion: 82.79<br>Late Fusion (a): 83.39<br>**Late Fusion (b): 83.76** |

**Table 4.** EDA performance comparison to early work on the BVDB (Part A) in a LOSO cross-validation setting. The performance metric consists of the average accuracy (in %) over the LOSO cross-validation. The best performing approach for each classification task is depicted in bold.

| Method | $T_0$ vs. $T_1$ | $T_0$ vs. $T_2$ | $T_0$ vs. $T_3$ | $T_0$ vs. $T_4$ |
| --- | --- | --- | --- | --- |
| Werner et al. [32] | 55.40 | 60.20 | 65.90 | 73.80 |
| Lopez-Martinez et al. [50] | 56.44 | 59.40 | 66.00 | 74.21 |
| **Our Approach (CNN)** | **61.67** | **66.72** | **76.06** | **85.03** |

**Fig 5.**
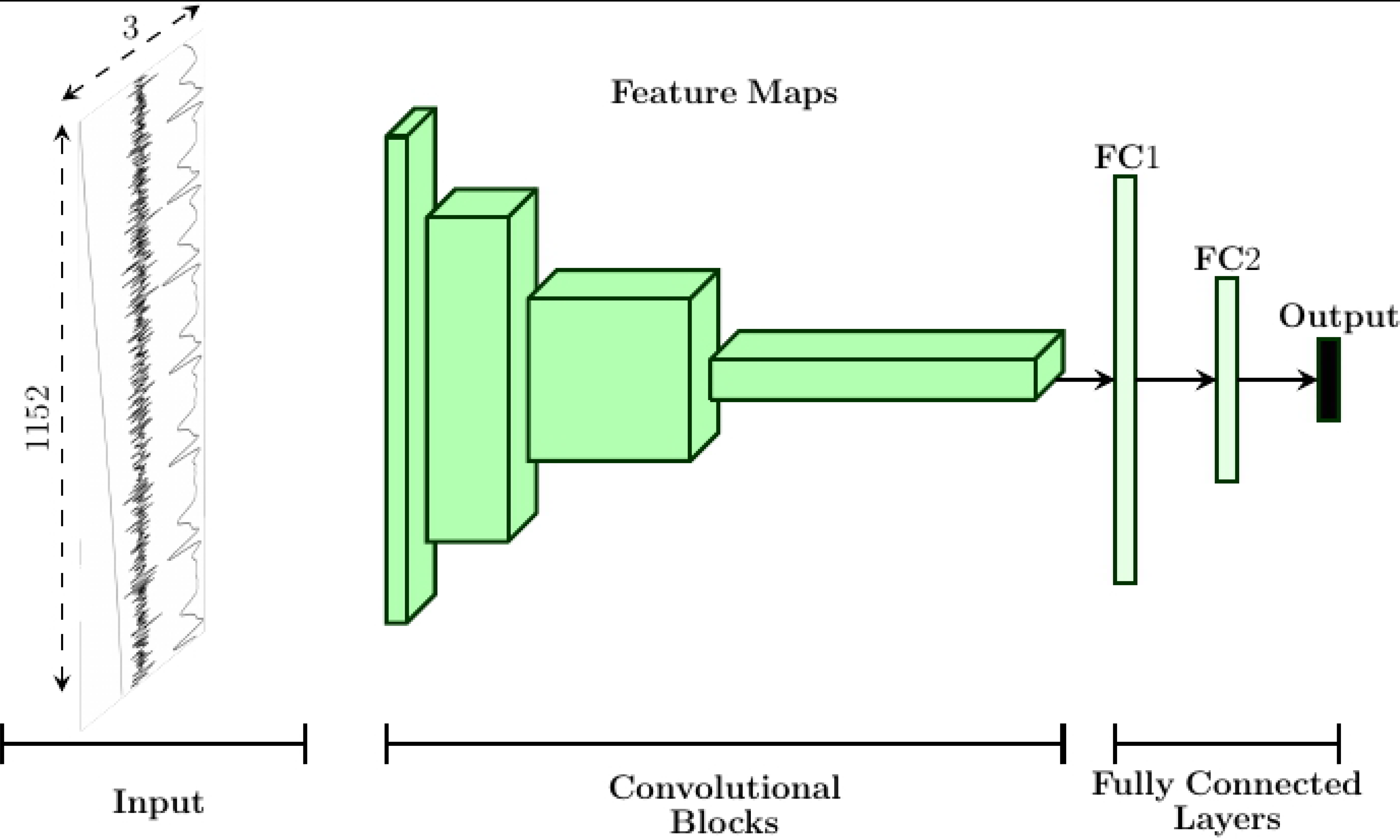
Late fusion parameters optimization. (a) Performance of each modality specific architecture, as well as of the late fusion approach, for each participant regarding the classification task *T*_0_*vs*.*T*_4_. The lower the performance, the darker the corresponding color. (b) Box plot of the weighting parameters *α*_1_, *α*_2_, *α*_3_ computed during the LOSO cross-validation evaluation of the classification task *T*_0_*vs*.*T*_4_. Within each box plot, the mean and median weight values of the performed LOSO cross-validation evaluation are respectively depicted with a dot and a horizontal line.

Finally, further experiments including the discrimination between the lowest temperatures of elicitation and the highest temperature of elicitation, as well as a multi-class experiment involving all levels of pain elicitation are conducted. In Fig 6, an overview of the classification results for both binary classification experiments *T*_0_*vs*.*T*_4_ and *T*_1_*vs*.*T*_4_, as well as for the multi-class classification experiment (*T*_0_*vs*.*T*_1_*vs*.*T*_2_*vs*.*T*_3_*vs*.*T*_4_) are depicted. During these experiments, only the second late fusion architecture was used to perform the aggregation of the information stemming from all the physiological modalities. Regarding the classification task *T*_1_*vs*.*T*_4_, the late fusion approach performs slightly better than EDA with an average accuracy of 76.81% (compared to an average accuracy of 76.70% for EDA), however not significantly. Moreover, the performances of the models specific to both ECG and EMG in the case of a 5-class classification experiment (see Fig 6(c)) are slightly above chance level (average accuracy of 23.30% and 23.17% for ECG and EMG respectively), while the performance of EDA is significantly above chance level with an average accuracy of 37.15%. Thus, the late fusion approach of all modalities is unable to outperform EDA with an average accuracy of 36.26%, since most of the models’ outputs specific to ECG and EMG are mostly unreliable. These results also suggest that the discrimination between all the 5 levels of pain elicitation is a very difficult classification task.

**Fig 6.**
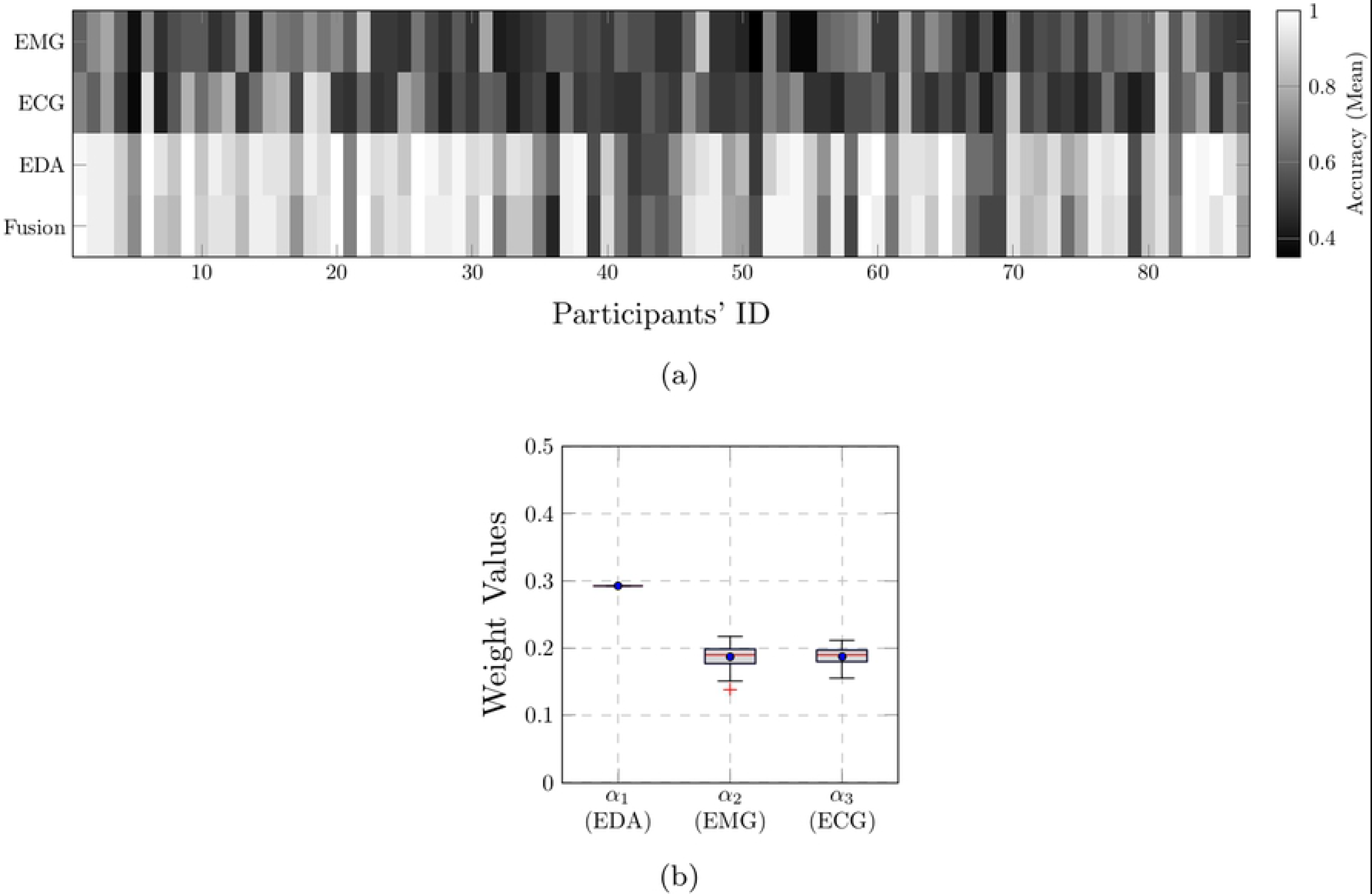
Classification results. (a) *T*_0_*vs*.*T*_4_. (b) *T*_1_*vs*.*T*_4_. (c) *T*_0_*vs*.*T*_1_*vs*.*T*_2_*vs*.*T*_3_*vs*.*T*_4_. The performance of a random classifier in the case of a 5-class classification task is 20%. Within each box plot, the mean and median classification accuracy across all 87 participants are depicted respectively with a dot and a horizontal line.

## Discussion and Conclusion

This work explored the application of deep neural networks for pain intensity classification based on physiological data including ECG, EMG and EDA. Several CNN architectures, based on 1-D and 2-D convolutional layers, were designed and assessed based on the *BioVid Heat Pain Database (Part A)*. Furthermore, several deep fusion architectures were also proposed for the aggregation of relevant information stemming from all involved physiological modalities. The proposed architecture specific to EDA significantly outperformed the results presented in previous works in all classification settings. For the classification task *T*_0_*vs*.*T*_4_, EDA achieved a state-of-the-art average accuracy of 85.03%. The proposed late fusion approach based on a weighted average of each modality specific model’s output also achieved state-of-the-art performances (average accuracy of 83.76% for the classification task *T*_0_*vs*.*T*_4_), but was unable to significantly outperform the deep model based uniquely on EDA. It is believed that a better optimization of the weighting parameters could potentially improve the current performance of the fusion architecture.

Moreover, all designed architectures were trained in an *end*-*to*-*end* manner. Therefore, it is also believed that pre-training and fine tuning at different levels of abstraction of the CNN architectures, as well as the combination with recurrent neural networks (in order to include the temporal aspect of the physiological signals in the inference model), could also potentially improve the performance of the current system, since such approaches have been successfully applied in other domains of application such as facial expression recognition [51–53]. Finally, the recorded video data provides an additional channel that can be integrated into the fusion architecture in order to improve the performance of the whole system. Therefore, the video modality should also be evaluated and assessed in combination with the physiological modalities.

In summary, the performed assessment suggests that deep learning approaches are relevant for the inference of pain intensity based on 1-D physiological data, and such methods are able to significantly outperform traditional approaches based on hand-crafted features. Domain expert knowledge could be bypassed by enabling the designed deep architecture to learn relevant features from the data. In the future iterations of the current work, approaches consisting of combining both learned and hand-crafted features should be addressed. Also, the designed architectures should be also assessed by replacing the classification experiments by regression experiments. Additionally, several data transformation approaches applied to the recorded 1-D physiological data in order to generate 2-D visual representations (e.g. spectrograms) should also be assessed in combination with established deep neural network approaches, specifically designed for this type of data representation.

## Acknowledgments

The research leading to these results has received funding from the Federal Ministry of Education and Research (BMBF, SenseEmotion) to FS, (BMBF, e:Med, CONFIRM, ID 01ZX1708C) to HAK, and the Ministry of Science and Education Baden-Württemberg (Project ZIV) to HAK. Peter Bellmann is supported by a scholarship of the Landesgraduiertenföorderung Baden-Württemberg at Ulm University. We gratefully acknowledge the support of NVIDIA Corporation with the donation of the Tesla K40 GPU used for this research.

